# Administration of vaccine-boosted COVID-19 convalescent plasma to SARS-CoV-2 infected hamsters decreases virus replication in lungs and hastens resolution of the infection despite transiently enhancing disease and lung pathology

**DOI:** 10.1101/2023.08.22.553458

**Authors:** Timothy D. Carroll, Talia Wong, Mary Kate Morris, Clara Di Germanio, Zhong-min Ma, Mars Stone, Erin Ball, Linda Fritts, Arjun Rustagi, Graham Simmons, Michael Busch, Christopher J. Miller

## Abstract

The utility of COVID-19 convalescent plasma (CCP) for treatment of immunocompromised patients who are not able to mount a protective antibody response against SARS-CoV-2 and who have contraindications or adverse effects from currently available antivirals remains unclear. To better understand the mechanism of protection in CCP, we studied viral replication and disease progression in SARS-CoV-2 infected hamsters treated with CCP plasma obtained from recovered COVID patients that had also been vaccinated with an mRNA vaccine, hereafter referred to as Vaxplas. We found that Vaxplas dramatically reduced virus replication in the lungs and improved infection outcome in SARS-CoV-2 infected hamsters. However, we also found that Vaxplas transiently enhanced disease severity and lung pathology in treated animals likely due to the deposition of immune complexes, activation of complement and recruitment of increased numbers of macrophages with an M1 proinflammatory phenotype into the lung parenchyma.

## Introduction

Beginning in fall 2019 in China a novel respiratory disease, COVID-19, was seen in a subset of people infected with Sudden Acute Respiratory Syndrome Coronavirus-2 (SARS-CoV-2) [1]. Due to the historical success of convalescent plasma in treatment of other viral diseases, infusion of COVID-19 convalescent plasma (CCP) obtained from recovered COVID-19 patients was tested early in the pandemic [2–4]. CCP with high titer neutralizing antibodies, administered within 72 hours of symptom onset, decreases disease progression, hospitalization, and mortality [5,6]. While anti-spike monoclonal-antibodies were used to successfully treat COVID-19 early in the pandemic, more transmissible, monoclonal antibody-resistant SARS-CoV-2 variants have emerged as SARS-CoV-2 has evolved [7–12]. In contrast, CCP obtained from patients recovering from currently circulating SARS-CoV-2 variants maintains clinical efficacy against emerging SARS-CoV-2 variants [13,14]. In January 2022, the FDA revised the EUA of CCP to include patients who are hospitalized with impaired humoral immunity [15]. In a randomized trial, while patients in the CCP arm tended to exhibit worsening pulmonary function at day 4 post-infusion, this did not abrogate the positive impact of CCP on survival of immunosuppressed patients [16].

To understand CCP’s mechanism of protection, we studied viral replication and disease progression in SARS-CoV-2 infected hamsters treated with CCP obtained from recovered patients subsequently vaccinated with an mRNA vaccine, hereafter referred to as Vaxplas. We found that Vaxplas dramatically reduced SARS-CoV-2 replication in the lungs and improved clinical outcome.

## Results

### Human IgG and anti-spike binding antibody levels in hamsters infused with human plasma

Forty male 8–10-week-old Syrian Golden hamsters were intranasally inoculated with 5000 PFU of a mixture of seven SARS-CoV-2 strains (Table 1). Twenty-four hours after the virus inoculation, 1 group of 10 hamsters was infused with 2 ml of Vaxplas (Table 2), a second group of 10 hamsters was infused with 2 ml of CCP from unvaccinated, convalescent COVID patients (Table 2), a third group of 10 hamsters was infused with 2 ml of control plasma (CoP) from SARS-CoV-2 naïve human donors (Table 2) and a fourth group 10 hamsters was untreated. All infusions were performed by peritoneal injection. To assess lung histology and viral loads, five animals from each group were necropsied at day 3 PI, and the remaining animals were necropsied at day 6 PI.

**Table 1:**
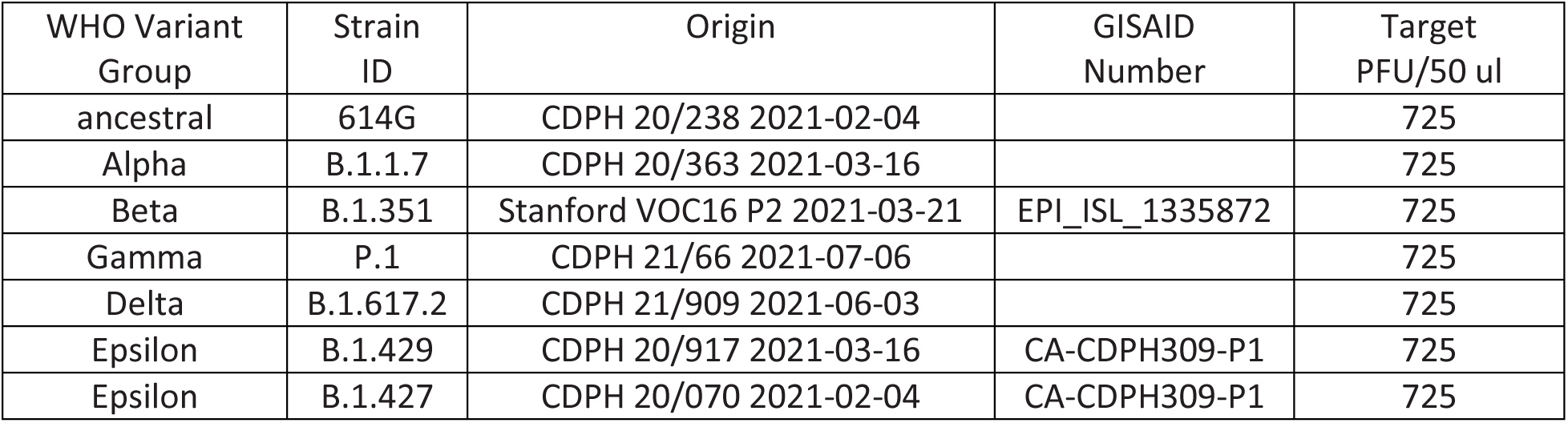
SARS-CoV-2 variants used to produce the mixed inoculum used in these studies.

| WHO Variant Group | Strain ID | Origin | GISAID Number | Target PFU/50 ul |
| --- | --- | --- | --- | --- |
| ancestral | 614G | CDPH 20/238 2021-02-04 |  | 725 |
| Alpha | B.1.1.7 | CDPH 20/363 2021-03-16 |  | 725 |
| Beta | B.1.351 | Stanford VOC16 P2 2021-03-21 | EPI_ISL_1335872 | 725 |
| Gamma | P.1 | CDPH 21/66 2021-07-06 |  | 725 |
| Delta | B.1.617.2 | CDPH 21/909 2021-06-03 |  | 725 |
| Epsilon | B.1.429 | CDPH 20/917 2021-03-16 | CA-CDPH309-P1 | 725 |
| Epsilon | B.1.427 | CDPH 20/070 2021-02-04 | CA-CDPH309-P1 | 725 |

**Table 2:** Human plasma pools used in these studies.

| Human Plasma Pools | RVPN NT<sub>50</sub> | Dose and route of administration |
| --- | --- | --- |
| Human COVID-19 Convalescent (CCP) | 1049 | 2 ml Intraperitoneal |
| Human COVID-19 Convalescent, Vaccine Boosted (Vaxplas) | >14,000 | 2 ml Intraperitoneal |
| Human Pre-COVID-19 normal control (CoP) | Negative | 2 ml Intraperitoneal |

Two days after the infusions (day 3 PI) of the Vaxplas, CCP and CoP animals, h-IgG levels in plasma exceeded 50 ug/ml (range 74-197 ug/ml) (Figure 1A). As expected, h-IgG was undetectable in the virus only animals (Figure 1A). Five days after the infusions (day 6 PI), human IgG levels had declined (range 14-117 ug/ml) in plasma of the Vaxplas, CCP and CoP hamsters (Figure 1A). Human IgG was undetectable in the virus only animals at day 6 PI (Figure 1A). These results demonstrate the successful transfer of human IgG to the treated animals.

**Figure 1.**
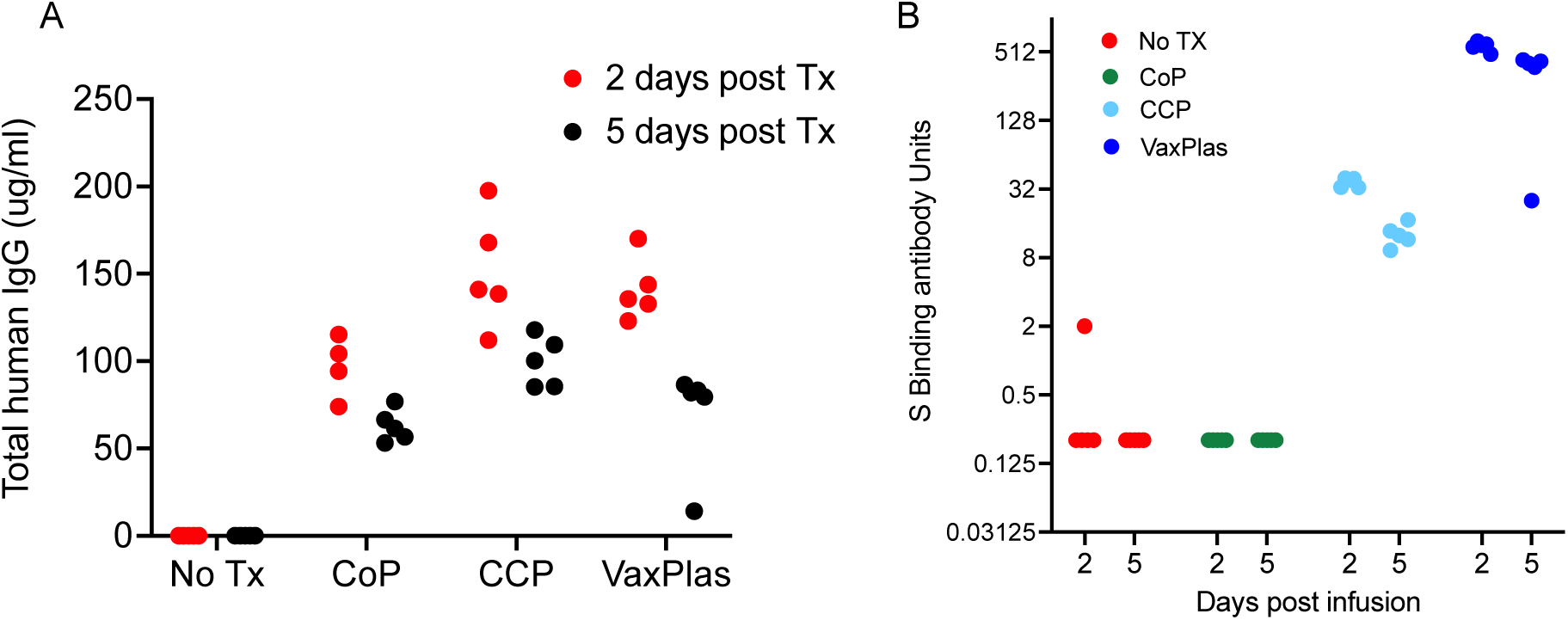
Human IgG levels and spike binding antibody levels in hamsters infused with human plasma after SARS-CoV-2 infection. A) Levels of human IgG in the blood of hamsters collected 2 and 5 days after intraperitoneal infusion of Vaxplas, CCP and CoP. B) Levels of anti-spike antibodies in the blood of hamsters collected 2 and 5 days after plasma infusion.

Two days after the infusions (day 3 PI), anti-S antibody levels were high (>512 S binding antibody units [BAU]) in the Vaxplas group, moderate (30-40 S BAU/mL) in the CCP group, but generally undetectable in the CoP and virus only group; one virus-only animals had very low levels of S antibodies (2 S BAU/mL) that may represent a nascent IgM response of this hamster to the infection (Figure 1B). Five days after the infusions (day 6 PI), anti-S antibody levels remained moderate to high (range 32-512 S BAU/mL) in the Vaxplas group, low (9-17 BAU/mL) in the CCP group, and undetectable in the CoP and virus only group (Figure 1B). Thus, anti-SARS-CoV-2 spike IgG binding antibodies were transferred by the infusions, and the Vaxplas animals had at least 10-fold higher anti-S IgG antibody levels than the CCP group.

### Effect of CCP on Disease Severity

We used change in body weight to assess disease severity in the infected animals. Body weight transiently declined in all the hamsters inoculated with SARS-CoV-2. The greatest body weight declines occurred in the virus only and CoP treated animals at day 6 PI. Body weights were lowest in CCP treated animals at day 2 PI and at day 3 in the Vaxplas treated animals but did not decline further at day 6 (Figure 2A). At day 3 PI the differences in body weight change (BWC) between the CoP, CCP and Vaxplas groups were significant, with the CoP animals having more severe weight loss than the CCP animals at day 3 PI (Figure 2B). In addition, at day 3 PI, the Vaxplas animals had more severe weight loss than the CCP animals (Figure 2C). At days 6 PI, the extent of the BWC in the virus only and CoP groups were similar and significantly more marked than in the CCP and Vaxplas groups (Figure 2D). These results confirm that the CCP and Vaxplas treated animals had less severe disease than the untreated or CoP treated hamsters at day 6 PI. Unexpectedly, the CCP group had significantly more BWC than the CoP group at day 3 PI and the Vaxplas treated animals had significantly more BWC than the CCP group at day 3 PI, a difference that reached significance in the case of the CoP animals.

**Figure 2.**
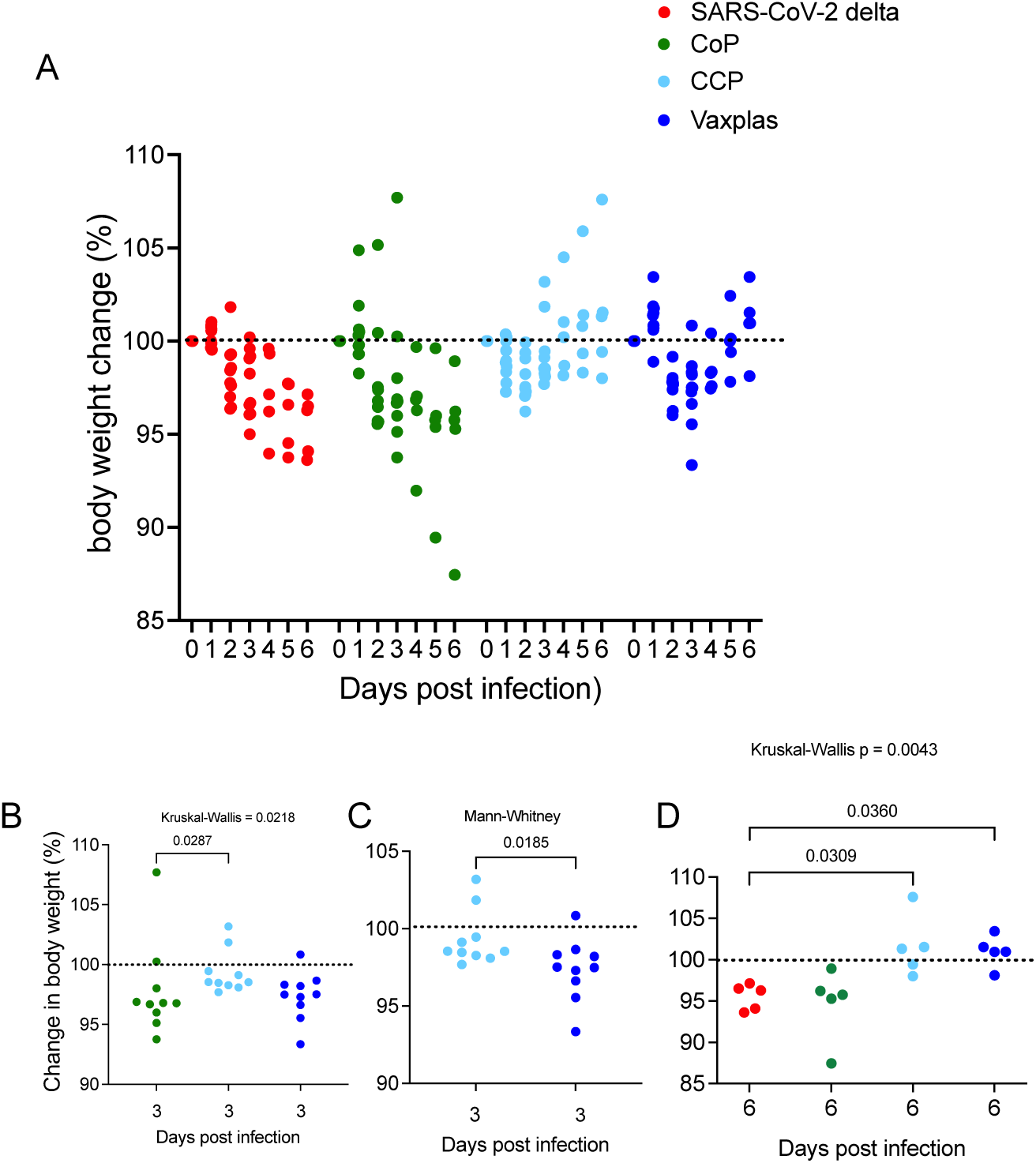
Percent body weight change in hamsters infused with human plasma after SARS-CoV-2 infection. A) Comparison of daily change in body weight (%) in treated and virus-only hamsters after SARS-CoV-2 infection. B) Comparison of BWC in treated hamsters 3 days after SARS-CoV-2 infection. C) Comparison of BWC in CCP and Vaxplas treated hamsters 3 days after SARS-CoV-2 infection. D) Comparison of BWC in treated hamsters 6 days after SARS-CoV-2 infection. The Kruskal-Wallis’s test with pairwise comparisons; panels B, and D, Mann-Whitney test; panel C.

### Effect of CCP on Virus Replication

At day 3 PI, the levels of sg RNA in the upper respiratory tract (URT) were very high (> 10^6^ copies/ug total RNA) in all 4 animal groups (Figure 3A). At day 6 PI, sg RNA levels in the URT were lower (generally < 10^6^ copies/ug total RNA) and more variable in all 4 animal groups (Figure 3B). Thus, CCP and Vaxplas had minimal effect on virus replication in the URT.

**Figure 3.**
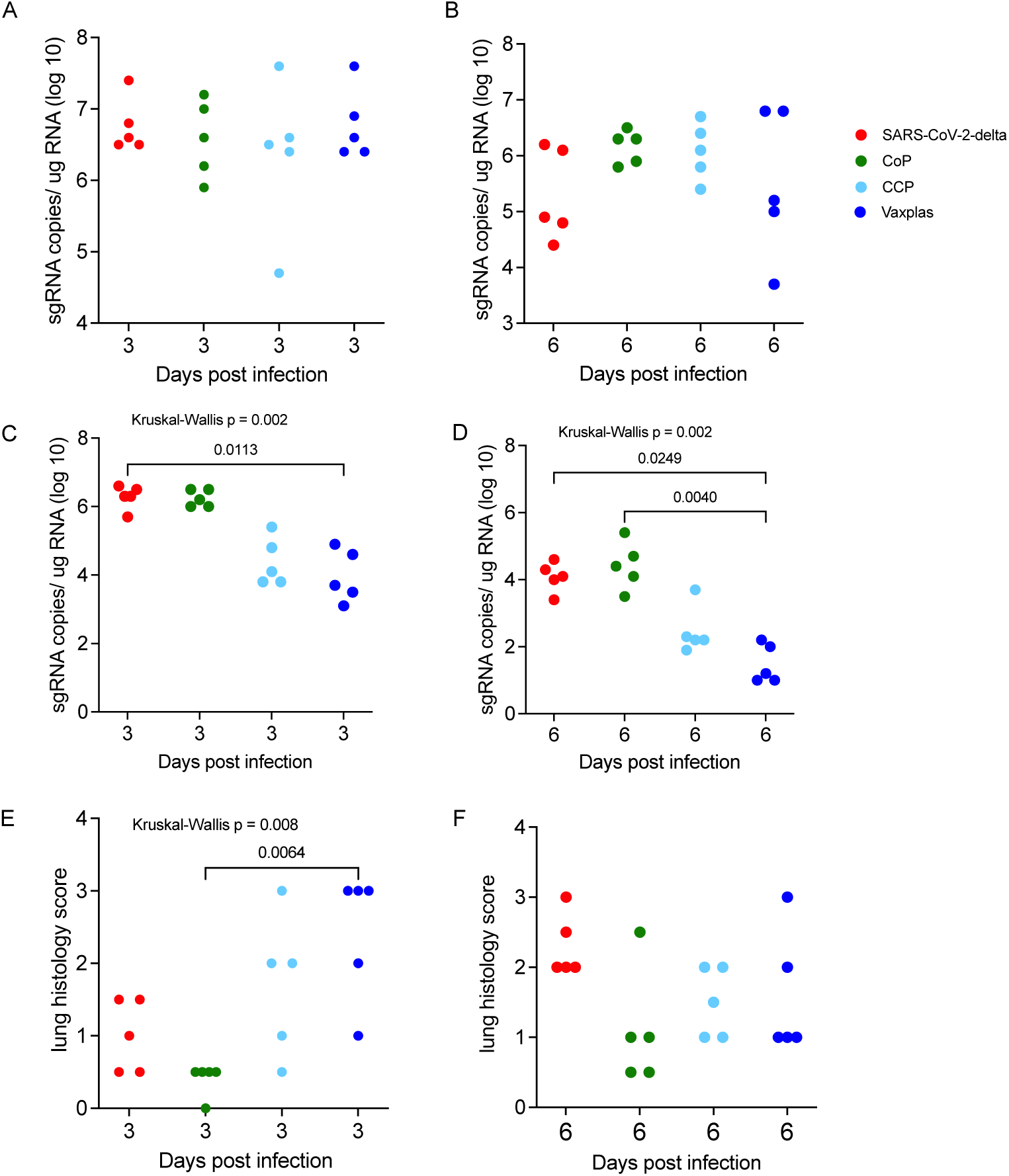
Virology and lung histopathology scores in hamsters infused with human plasma after SARS-CoV-2 infection. Comparison of sgRNA levels in A) URT of treated and control hamsters 3 days PI. B) URT of treated and control hamsters 6 days PI. C) lungs of treated and control hamsters 3 days PI. D) in lungs of treated and naïve control hamsters 6 days PI. Comparison lung histopathology scores of treated and control hamsters at E) 3 days PI and F) 6 days PI. Kruskal-Wallis’s test with pairwise comparisons; all panels.

At day 3 PI, the virus only and CCP animals had high levels of sgRNA in lungs (Figure 3C), while the CCP and Vaxplas treated animals had lower sgRNA in lungs compared to the mean levels of all 4 groups (Figure 2C). Pairwise comparison demonstrated a significantly lower sgRNA in lungs of Vaxplas-treated animals compared to virus only animals (Figure 3C). At day 6 PI, sgRNA levels were significantly different in the lungs of the 4 animal groups (Figure 3D). Pairwise comparisons demonstrated sgRNA were significantly lower in lungs of Vaxplas-treated animals compared to CoP animals and virus-only animals (Figure 3D). Thus, Vaxplas blunts SARS-CoV-2 replication in the lungs but not the URT of infected hamsters. CCP also lowers virus replication in the lungs but not to the same extent as Vaxplas. Our original rationale for using a mixed inoculum of 7 SARS-CoV-2 variants was to determine if replication of a particular variant in the mixture was more resistant to immune control by CCP and Vaxplas than other variants. However technical issues with the Quills assay [17] has delayed that analysis indefinitely.

### Effect of CCP on Lung Pathology

The lungs of the animals were evaluated histologically, and the total area of diseased lung was estimated and scored. In the virus only and CoP animal groups, lung lesions were less extensive at 3 days PI and more extensive at 6 days PI (Fig 3E and 3F), while in the CCP and Vaxplas groups lung lesions were more extensive at 3 days PI and less extensive at 6 days PI (Fig 3E and 3F). At day 3 PI the differences in the extent of lung disease among the groups were significant, and pairwise comparison demonstrated that there was significantly more disease in the Vaxplas group compared to the CoP group (Figure 3E). At day 6 PI, lung disease was most widespread in the virus only group, lowest in the CoP group and intermediate in the CCP and Vaxplas groups, although the differences among the groups were not significant (Figure 3F).

The nature of the lesions in the virus only, CoP and CCP treated animals were similar and consistent with previous reports [17,18]. At day 3 PI, these 3 groups had moderate multifocal necro-suppurative bronchiolitis (Fig 4 A, B) with perivascular lymphocytic cuffing and endotheliitis in small arteries. Variable bronchiolar epithelial hyperplasia and scattered type II pneumocyte hyperplasia were also noted. The changes in the lungs of the Vaxplas animals at day 3 PI (loss of alveolar septal architecture, hemorrhage, edema, fibrin, necrotic debris and mixed inflammation, perivascular cuffing and endotheliitis) were more severe and extensive than in the other groups (Figure 3C and 4C). At day 6 PI, animals in all groups had necrotizing, neutrophilic and histiocytic bronchointerstitial pneumonia with syncytial cells, perivascular cuffing, and endotheliitis, with prevalent bronchiolar epithelial hyperplasia and type II pneumocyte hyperplasia (Fig 4 D, E, F). However, the inflammation and hemorrhage in the Vaxplas animals had resolved to a greater extent than the other groups.

**Figure 4.**
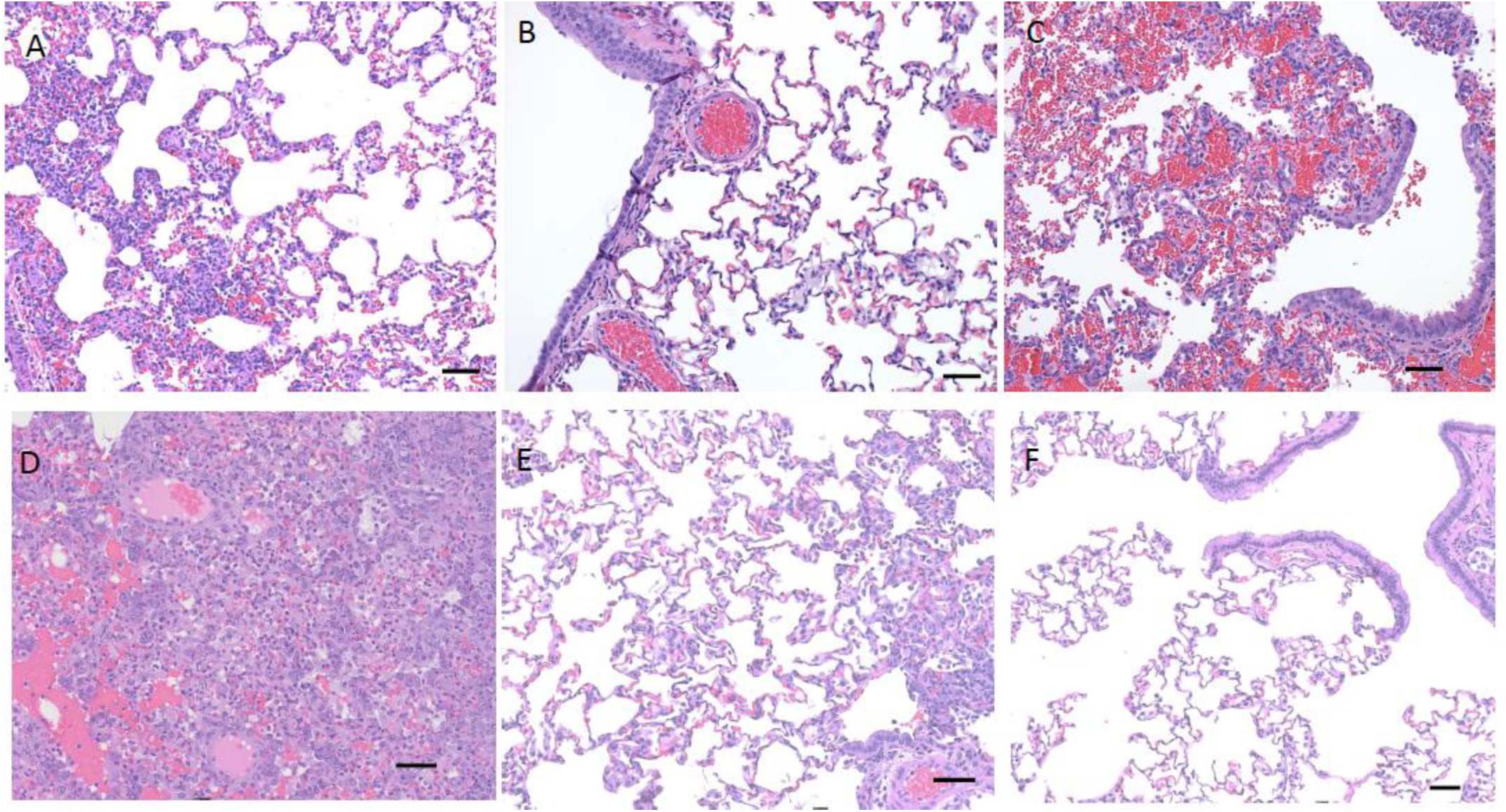
Histopathology of lungs of hamsters infused with human plasma after SARS-CoV-2 infection. Day 3 PI, A-C. A) Virus-only animals, B) CoP animals and C) Vaxplas animals. Day 6 PI, D-F. D) Virus-only animals, E) CoP animals and F) Vaxplas animals. IBA1+ cells in G) Virus-only animals, H) CoP animals and I) Vaxplas animals. Hematoxylin and Eosin stain. Scale bars = 50 microns

In day 3 PI lung sections of virus-only animals, human (h)-IgG was undetectable but the C3 complement fragment was found in the lumen and walls of small and medium blood vessels and capillaries within alveolar septa (Fig 5 A, D). In the CCP and Vaxplas animals, h-IgG and C3 were localized in the medium size blood vessels and in alveolar septa capillaries (Fig 5 B, C, E, F). In the CoP animals, h-IgG and C3 were found only in the lumen and walls of medium and small blood vessels with little staining in alveolar septa capillaries (Fig 5 B, E). In the Vaxplas animals, extravascular h-IgG and C3 were also found in the alveolar spaces and inflamed areas of the pulmonary parenchyma (Fig 5 C, F).

**Figure 5.**
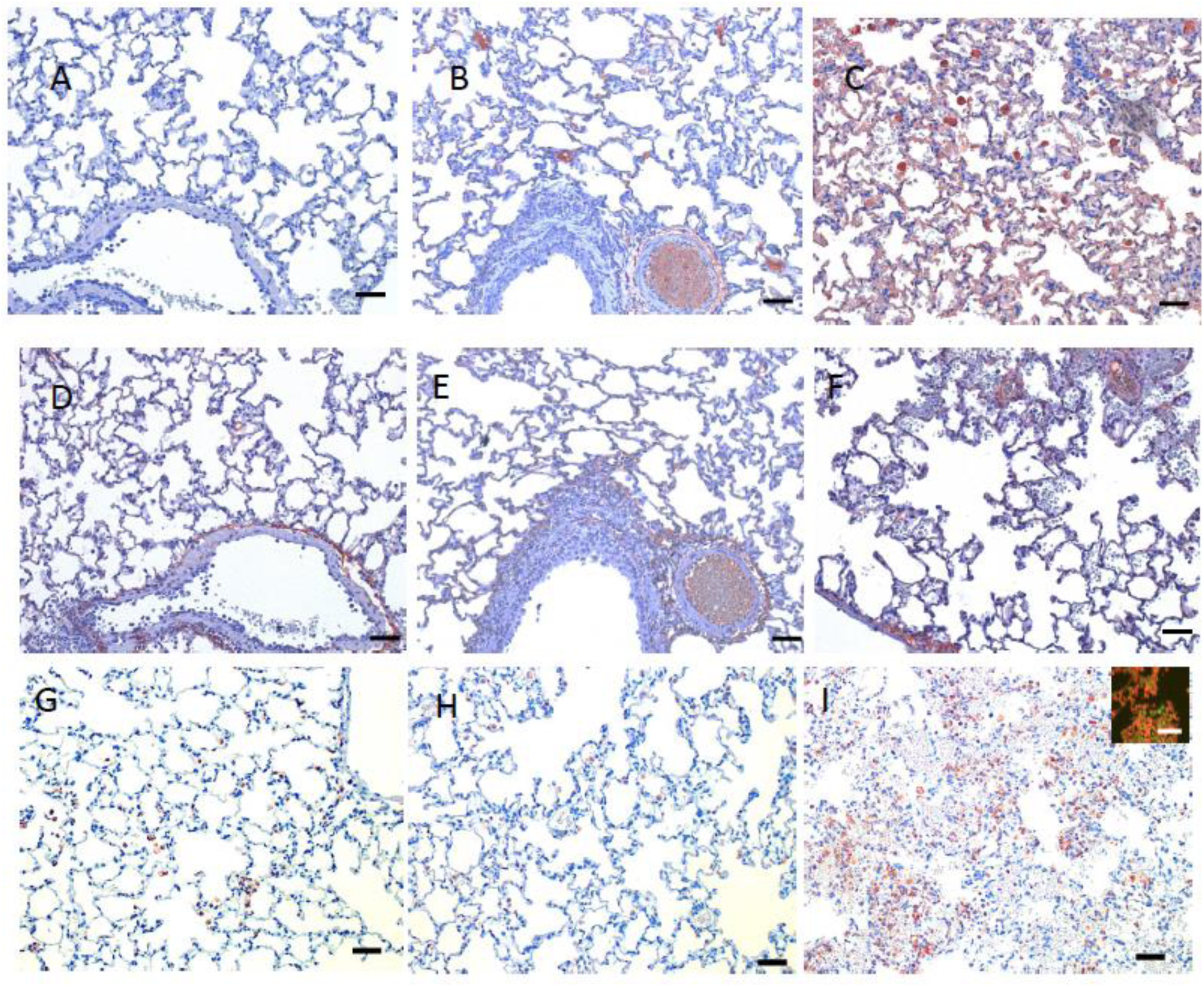
Immunohistochemical staining for human IgG, C3 fragment of complement and IBA1+ macrophages in lungs of hamsters infused with human plasma after SARS-CoV-2 infection. Human IgG staining A) Virus-only animals, B) CoP animals, C) Vaxplas animals. C3 staining D) Virus-only animals, E) CoP animals F) Vaxplas animals. IBA1+ cells G) Virus-only animals, H) CoP animals and I) Vaxplas animals, inset: IBA1 and Stat-1 double immunofluorescent staining of macrophages. Hematoxylin counterstain. Scale bars = 50 microns

In all the lungs of all animals, variable numbers of IBA1+ macrophages were found in and around inflamed airways and blood vessels, and in inflamed alveolar septa and alveolar spaces (Fig 5 G, H, I). The number of IBA1+ macrophages was highest in the lungs of the Vaxplas (Fig 5 I), moderate in the CCP animals and the virus-only animals (Fig 5 G) and lowest in the CoP animals (Fig 5 H). Double label immune fluorescent staining demonstrated that > 90% of IBA1+ macrophages were Stat 1+ (inset, Fig 5 I), indicating that most macrophages in the lungs of SARS-CoV-2 infected hamsters, regardless of treatment, had a pro-inflammatory M1 phenotype. In fact, the densities of IBA-1+ [19] and Stat 1+ [20–22] macrophages were significantly higher in the Vaxplas vs the CoP animals (Fig 6 A, B).

**Figure 6.**
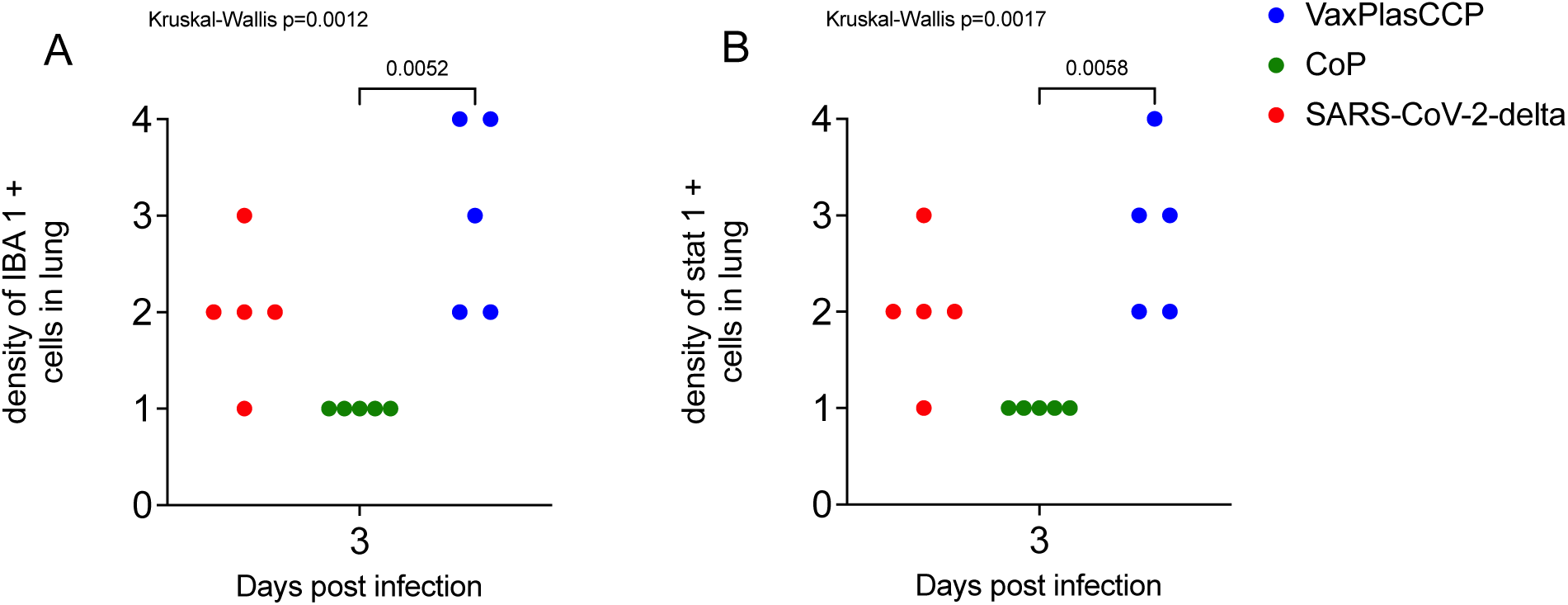
Density of IBA1+ macrophages and Stat-1+ cells in lungs of hamsters infused with human plasma after SARS-CoV-2 infection. A) Comparison IBA1+ macrophages in lungs of treated and virus-only hamsters 3 days after SARS-CoV-2 infection. B) Comparison Stat-1 + cells in lungs of treated and virus-only hamsters 3 days after SARS-CoV-2 infection. Kruskal-Wallis’s test with pairwise comparisons; both panels.

## Discussion

We found that Vaxplas improved disease outcome and dramatically reduced virus replication in the lung, but not the URT, of SARS-CoV-2 infected hamsters. Peritoneal infusion of Vaxplas, CCP and CoP resulted in transfer of human IgG to the hamsters in the range of 50-200 ug/ml (0.5-20 mg/dl) hamster plasma, a concentration that is about 50-100 fold lower than the median IgG levels found in adult humans [23]. Thus, while the plasma infusions successfully transferred human IgG to the hamsters, the dose of IgG that the animals received was modest. The infusion of CCP and Vaxpals resulted in transfer of low (CCP) to moderate titers (Vaxplas) of anti-S antibodies to the hamsters. For comparison, anti-S antibody titers 4 months after vaccination of previously infected people had a median of 19,539 BAU/mL [24]. Further, one recent metanalysis of 5 published CCP transfusion studies found that the virus neutralizing antibody titers in the CCP units used ranged from 8 to 14,580 as determined in several different assays and labs [25]. Although, post transfusion titers have not determined in human recipients of CCP, the neutralizing antibody titers in the plasma of infected humans and CCP units are up to 40 times higher than the highest titers (512 S BAU/mL) found in the plasma of Vaxplas hamsters and 600 times higher than the highest plasma titers found in the CCP hamsters (40 S BAU/mL). The extent of the reduction in viral replication in these VAXplas and CCP groups was consistent with the relative amount of anti-S antibodies transferred to the 2 groups of hamsters. (median anti-S antibody titers in the Vaxplas group were approximately 16 times higher than in the CCP group and the median sgRNA levels were 5-10 x lower than in the Vaxplas group compared to the CCP group.

Despite the modest levels of anti-S antibodies transferred to the hamsters, the infusions improved the course of SARS-CoV-2 disease. By the end of the study (6 days PI) both the CCP animals and the Vaxplas animals had significantly less body weight loss than the virus only animals and there was a similar trend with the CoP group. While this milder disease course was apparent in the minimal body weight loss of CCP animals relative to CoP animals at day 3 PI Figure 1B), it was not apparent in the Vaxplas animals. In fact, the body weight loss in the Vaxplas animals was significantly greater than in the CCP animals at day 3 PI, indicating that Vaxplas animals had a transient enhancement of disease severity that was not seen in the CCP group. This enhanced disease occurred even though the Vaxplas animals had the lowest lung viral loads of any animal group at day 3 PI.

Although lung sgRNA levels significantly lower than the virus-only group at day 3 PI, we also found that Vaxplas transiently enhanced disease severity and lung pathology, likely due to the deposition of immune complexes, activation of complement, and recruitment of macrophages with an M1 proinflammatory phenotype into the lung parenchyma. These results may explain why some patients receiving CCP transiently exhibit decreased pulmonary function around day 4 post-infusion [16].

Despite lung sgRNA levels as high or higher than the virus-only group, the CoP group had the lowest lung pathology scores at day 3 and day 6 PI and fewest lung IBA1+ macrophages and least C3 deposition in lung parenchyma at day 3 PI. Thus, it appears that normal human plasma containing no anti-SARS-CoV-2 antibodies profoundly reduced lung disease in the SARS-CoV-2 infected hamsters, while having no effect on virus replication. Studies are underway attempting to understand the mechanism behind this observation with the hypothesis that the CoP mitigates the pulmonary endotheliopathy commonly seen in COVID-19 patients [26] and SARS-CoV-2 infected hamsters [17].

In this study we found that infusion of hamsters 24 hours after SARS-CoV-2 infection with moderate titer CCP and high titer Vaxplas blunts virus replication in the lungs and improves the course of viral disease. In addition, we found that although normal plasma has no effect on clinical disease (weight loss) or virus replication in hamsters, it does decrease the severity and extent of inflammation in the lungs.

## Materials and Methods

### Ethics statement

All animal experiments were approved by the Institutional Animal Care and Use Committee of UC Davis (Protocol # 22233) and performed following the guidelines and basic principles in the United States Public Health Service Policy on Humane Care and Use of Laboratory Animals and the Guide for the Care and Use of Laboratory Animals. The work with infectious SARS-CoV-2 under BSL3 conditions was approved by the UC Davis Institutional Biosafety Committee.

### Human Plasma

Plasma pools (Table 2) made from aliquots of frozen plasma collected from human donors were used. Details of the control pool of normal plasma containing no anti-SARS-CoV-2 antibodies and CCP pool have been provided in previously published studies [27,28]. The CCP pool had a 50% neutralization titer (NT50) of 1,149 in a reporter virus assay (23). The Vaxplas plasma pool was made by combining plasma aliquots from three individual donors that had been vaccinated after recovering from a documented SARS-CoV-2 infection to achieve a pool that had a 50% neutralization titer (NT50) of 9,901 in a reporter virus assay. Two of the 3 Vaxplas pool donors were immunized with 2 doses of the FDA EUA-approved, monovalent Moderna vaccine and 1 donor was immunized with 2 doses of the FDA EUA-approved, monovalent Pfizer vaccine. The interval between the second immunization and plasma donations was between 2 and 6 months.

### Hamsters

7-9 week old male Syrian Hamsters purchased from Charles River Inc. were used.

### Viruses and cells

SARS-CoV-2 variants were isolated from patient swabs by CDPH, Richmond CA and Stanford University, Palo Alto CA and sequence verified as previously described [17].

### Hamster inoculations

For experimental inoculations, hamsters were infected intranasally with a total dose of variant mixture containing approximately 5000 PFU of SARS-CoV-2 suspended in 50μL sterile DMEM as previously described [17]. The dose of all variants was approximately equal (<10-fold difference).

### qPCR for sub-genomic RNA quantitation

Quantitative real-time PCR assays were developed for detection of full-length genomic vRNA (gRNA), sub-genomic vRNA (sgRNA), and total vRNA as previously described [17].

### Histopathology

At necropsy, lungs were evaluated blindly by a board certified veterinary anatomic pathologist blindly and the area of inflamed tissue (visible to the naked eye at subgross magnification) was estimated as a percentage of the total surface area of the lung section. The scores for healthy normal hamsters were set at zero.

### Immunohistochemistry and immunofluorescence staining of lung sections

Rabbit monoclonal (EPR4421) anti-Human IgG (Abcam), Rabbit polyclonal anti-C3 / C3b (EnenTex), Mouse monoclonal (GT10312) anti-IBA1 (Invitrogen), and Rabbit monoclonal (Tyr701)(58D6) anti-Phospho-Stat1 (Cell Signaling) antibodies were used. Briefly, 5µm paraffin sections were subjected to an antigen retrieval step consisting of incubation in AR10 (Biogenex) for 2 minutes at 125^0^C in the Digital Decloaking Chamber (Biocare), followed by cooling to 90^0^C for 10 minutes. The EnVision detection system (Agilent) was used with AEC (Agilent) as the chromogen. Slides were counterstained with Gill’s hematoxylin I (StatLab). Primary antibodies were replaced by mouse or rabbit isotype control (Thermo Fisher) and run with each staining series as the negative controls. In the double immunofluorescent stains of IBA1 and Phospho-Stat1, Goat anti-mouse Alexa Fluor 488 and Goat anti-Rabbit Alexa Fluor 568 (Invitrogen) were used to detect bound primary antibodies.

### Quantitation of Human IgG by ELISA

Total human IgG levels were determined following the manufacturer’s instructions in the Human IgG Total ELISA Kit (Invitrogen).

### Quantitation of anti-spike antibody levels

Anti-spike IgG levels were measured on the Ortho VITROS^®^ platform (Otho Clinical Diagnostics, Raritan, NJ USA) at Vitalant Research Institute as described [29]. Samples were tested following the manufacturer’s instructions, except that samples’ results above the limit of quantitation of >200 BAU/ml were further tested at 1:20 dilution.

### Statistical analysis

As noted in figure legends, median values were compared using the non-parametric Kruskal-Wallis’s and. A post-hoc Dunn multiple comparison test or Mann-Whitney tests was used for pairwise comparisons of the groups. For all analyses, differences with a p value < 0.5 were considered significant. Graph Pad Prism 9.3.1 (San Diego, CA) installed on a MacBook Pro (Cupertino, CA) running Mac OS Monterey Version 12.1 was used for the analysis.

## Acknowledgements

This work was supported by Intramural funding from the Center for Immunology and Infectious Diseases, UC Davis (CIID-2020-2), Vitalant A22-2082 and NIH/R01-AI118590 to C.J. Miller. The funders had no role in study design, data collection and analysis, decision to publish, or preparation of the manuscript.

## Author Contributions

Conceptualization: TDC, MB, CJM

Data curation: CJM, TDC, TW, EB, MKM, CDG, MS, LF, GH

Formal analysis: CJM, TDC, TW, EB

Funding acquisition: CJM, MB

Investigation: CJM, TDC, TW, EB, MKM, AR, CDG, MS, LF, GH, ZM

Methodology: CJM, TDC, MKM, AR, ZM, MS

Project administration: CJM

Supervision: CJM, MB

Writing – original draft: CJM

Writing – review & editing: TDC, TW, MKM, CDG, ZM, MS. LF, AR, GS, MB, CJM

